# Anthropogenically induced shifts in sediment vegetation impact coastal benthic microbiomes

**DOI:** 10.64898/2026.01.07.698198

**Authors:** Borja Aldeguer-Riquelme, Meagan A. Beatty, Luis H. Orellana, Esther Rubio-Portillo, Rudolf Amann, Gabrielle Potocki-Veronese, Josefa Antón, Fernando Santos

## Abstract

Opportunistic seaweeds of the genus *Caulerpa* have benefited from human-driven environmental change to expand within coastal lagoons, frequently displacing native seagrasses. In the Mar Menor lagoon (southeastern Spain), *Caulerpa prolifera* has progressively expanded since the 1970s, coinciding with the decline of the native seagrass *Cymodocea nodosa*. Sediments colonized by *Ca. prolifera* are characterized by elevated sulfide concentrations, which are toxic to seagrasses. Because sulfide is microbially produced, these sediment communities likely shape the competition between *Ca. prolifera* and *Cy. nodosa*. However, their metabolic potential and activity remain poorly understood. Here, we examined the taxonomic composition and functional capabilities of sediment microbial communities using 12 paired metagenomes and metatranscriptomes from areas dominated by either *Ca. prolifera* or *Cy. nodosa. Caulerpa*-associated sediments displayed a higher potential for organic matter degradation and sulfate reduction. Several metagenome-assembled genomes (MAGs) encoded enzymes putatively involved in the degradation of pectin-like polysaccharides characteristic of the *Caulerpa* cell wall. Comparison of metagenomic and metatranscriptomic data further revealed that the most abundant MAGs were not necessarily the most transcriptionally active. This finding challenges the common assumption that ecological importance is primarily determined by abundance and highlights the potential contribution of less abundant microorganisms to biogeochemical cycling. Further, 97 of the 100 most highly expressed genes lacked functional annotation. Recombinant production and functional characterization of the protein encoded by the most highly expressed gene in the analyzed sediments revealed that it is an enzyme able to breakdown endogenous bacterial peptidoglycan. Collectively, these results underscore the importance of integrating metagenomics and metatranscriptomics with experimental validation to achieve a more comprehensive understanding of microbial functional capabilities, an integration that is still uncommon in the microbial ecology field.

## Introduction

Coastal marine lagoons are critical environments due to the relevant ecosystem services they provide such as food provisioning, shore protection, or nesting areas for fish and birds. From an ecological perspective, coastal lagoons are productive environments chosen by many animal species for feeding and breeding [1]. In addition, vegetation contributes to stabilizing sediments and protects the shoreline from the erosion caused by waves [2]. Humans also use coastal lagoons for a variety of economic and recreational purposes including, for example, fishing, tourism or water sports. However, given their relatively small size and (semi)confined structure, they are also very sensitive to the impact of anthropogenic activities, such as eutrophication, mining activities or the introduction of allochthonous species [3].

Allochthonous species can be introduced in coastal lagoons accidentally (e.g. by ships) or voluntary (e.g. for aquaculture). Furthermore, changes in water temperature and salinity associated with climate change also facilitate the introduction and spread of allochthonous species to new habitats [4, 5]. Once introduced, they can exert multiple impacts on the ecosystem by competing with the local vegetation for resources (e.g., nutrients, light, surface), spreading new pathogens, and, in the most extreme cases, becoming invasive species capable of causing a shift in the ecosystem’s biodiversity [6].

The green seaweed genus Caulerpa contains notorious invasive species, such as *Caulerpa taxifolia* and *Caulerpa prolifera*, that have gained significant attention after their introduction and spread in non-native habitats. *Ca. prolifera* is a natural seaweed of the Mediterranean Sea that has been able to rapidly spread in coastal lagoons of Portugal, Spain or the United States [7–9], displacing the local seagrasses. The Mar Menor lagoon, located in southeast of Spain, provides an example of the invasive potential of *Ca. prolifera*. The seagrass *Cymodocea nodosa* is the predominant native species in this water body. However, due to the widening of channels connecting the Mar Menor lagoon with the Mediterranean Sea to allow the transit of ships, salinity decreased, allowing *Ca. prolifera* to introduce and spread across the lagoon over the past five decades [10]. *Ca. prolifera*, and the genus Caulerpa in general, are able to modify the sediment biogeochemical conditions, increasing organic matter content, sulfate reduction rates and consequently the sulfide pools [11–13]. Sulfide is known to be toxic for seagrasses [14], and therefore, the increase in sulfides could be the microbially-mediated mechanism behind the negative relationship observed between *Ca. prolifera* and Cy. nodosa. An alternative hypothesis suggests that *Ca. prolifera* imposes a nutrient limitation that hinders the growth and germination of Cy. nodosa [15]. The influence of *Ca. prolifera* on Cy. nodosa, however, is still under debate with some authors suggesting competitive exclusion [10, 15], while others dispute this notion [16].

Through 16S rRNA gene amplicon sequencing, we have previously observed distinct microbial communities associated with each species in the sediments, with a higher abundance of sulfate-reducing bacteria in *Ca. prolifera* sediments compared to those from Cy. nodosa [17]. However, due to the inherent limitations of amplicon-based approaches, the functional potential of these communities remained unresolved, constraining our understanding of both the ecological impact of *Ca. prolifera* on the lagoon and the potential mechanistic processes underlying its interaction with Cy. nodosa. Nonetheless, given the higher concentrations of sulfide and organic matter in *Ca. prolifera* sediments, we hypothesize that their associated microbial communities are enriched in anaerobic taxa with a greater metabolic potential for organic matter degradation, compared to those in Cy. nodosa sediments. Here, in order to address these hypotheses, we leveraged comparative metagenomic and metatranscriptomic analyses at both the community and genome levels to elucidate the metabolic differences between microbes in sediments dominated by *Ca. prolifera* and Cy. nodosa in the Mar Menor lagoon. We observed that *Ca. prolifera*-associated microbial communities were enriched in carbohydrate-active enzymes (CAZymes) and sulfate-reducing bacteria compared to *Cy. nodosa*. We found that the most abundant genomes were not necessarily the most expressed ones, challenging the classical vision of the abundant and rare biosphere in biogeochemical cycling. At the gene level, highest expression corresponded largely to genes coding for hypothetical proteins. The most highly expressed gene in the analyzed sediments has been characterized using recombinant overexpression in *E. coli*, which indicated that it corresponded to a previously undescribed cell wall–degrading enzyme.

## Results and discussion

### Overall community structure

In order to provide more insights into the impact of *Ca. prolifera* and *Cy. nodosa* (hereinafter, *Caulerpa* and *Cymodocea*, respectively) on the Mar Menor, six sampling stations were selected across the lagoon, three colonized with *Cymodocea* and three with *Caulerpa* (Figure 1). Two sampling campaigns were carried out in March and September 2018, collecting a total of 12 sediment samples which were analyzed using both metagenomics and metatranscriptomics to assess community composition and activity, respectively. Note that these same samples were previously studied using physicochemical variable measurements, 16S rRNA gene metabarcoding and cell density [17]. Metagenomes ranged from 28 to 71.4 Gb, with metagenome coverage, as estimated by Nonpareil, between 40% and 63% (Table 1). These values fall below or at the recommended threshold (60%) for achieving representative metagenomes and high-quality assemblies [18]. Accordingly, we acknowledge that our metagenomes did not capture the full microbial diversity present in the sediment, leaving a fraction of genomes and genes below the limit of detection. Consequently, some of the results presented here could potentially differ if metagenomes with higher diversity coverage were analyzed (e.g., >90%), as recently demonstrated [19]. In terms of alpha diversity, statistically significant (t-test, p-value = 0.01629) higher values of diversity were observed in *Cymodocea* metagenomes compared to those of *Caulerpa*, based on Nonpareil diversity values.

**Table 1.**
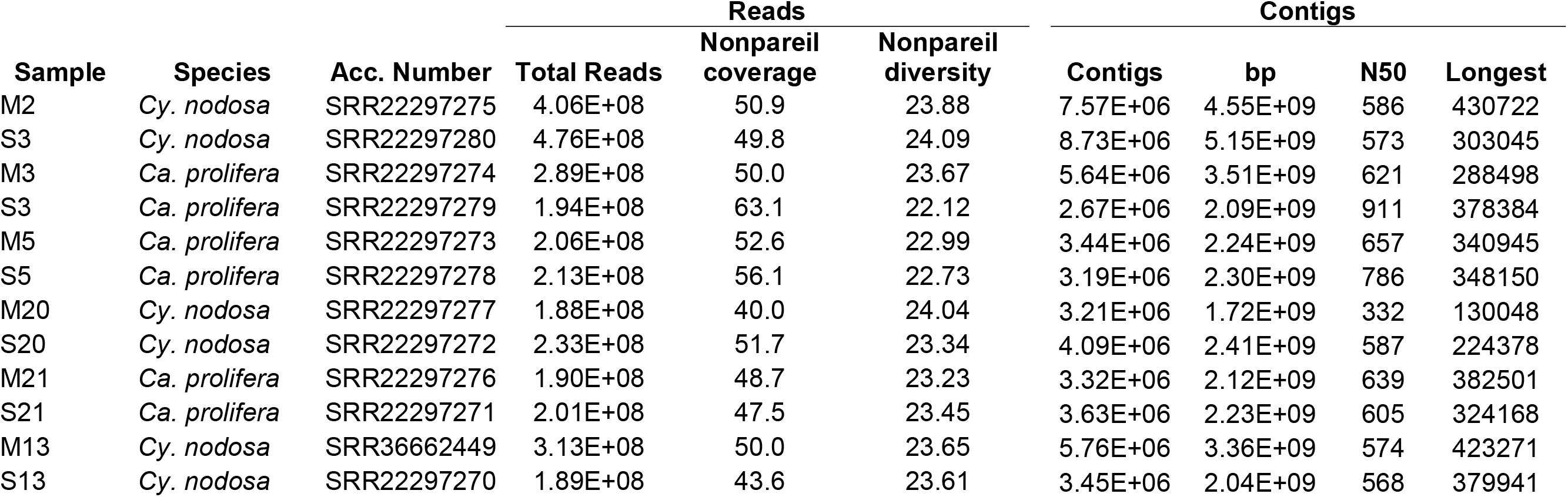
Characteristics of the 12 sediment metagenomes analysed in this study.

**Figure 1.**
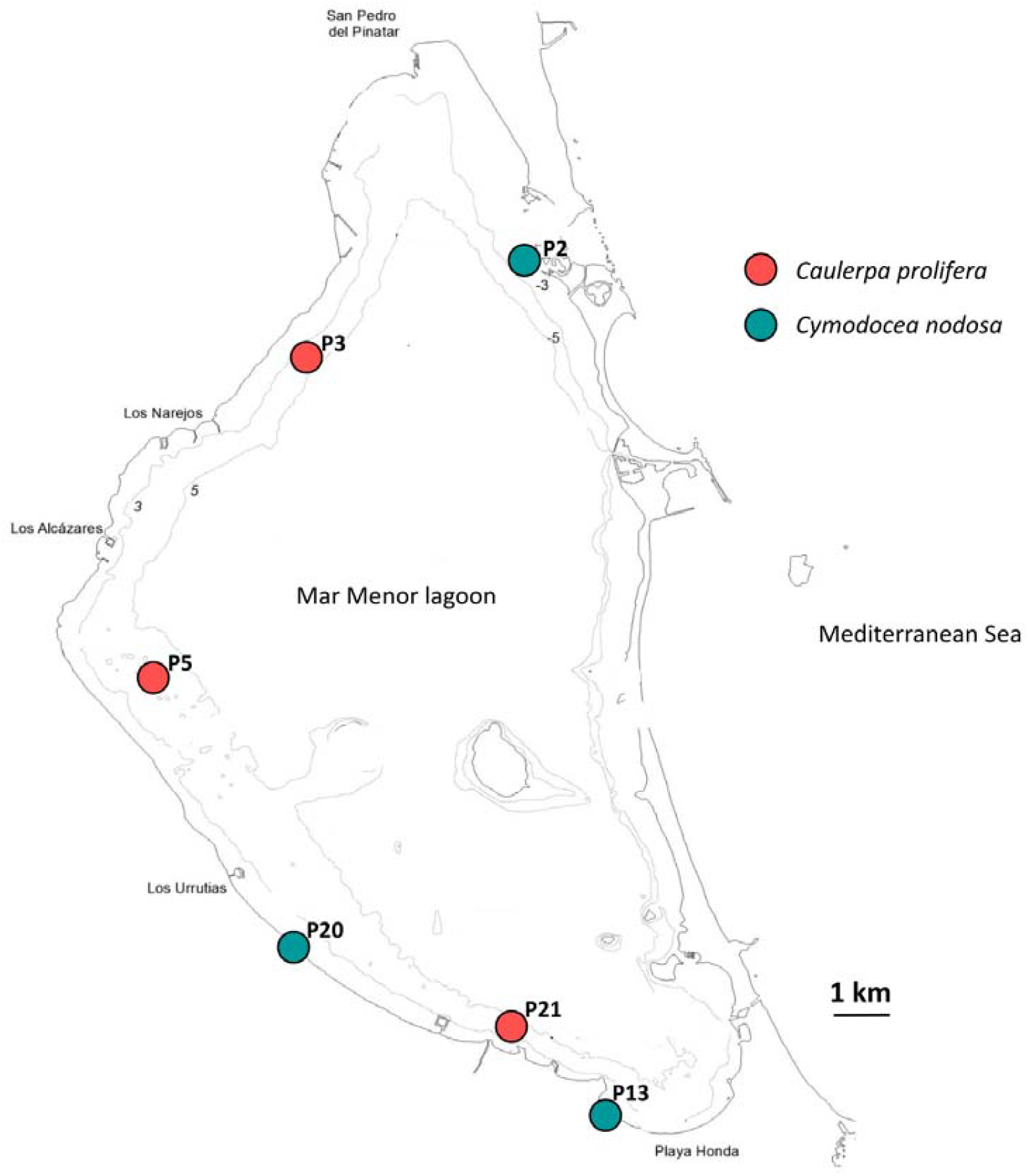
Sampling locations of sediment colonized by *Caulerpa prolifera* (red) and *Cymodocea nodosa* (blue) in the Mar Menor lagoon.

We first focused on the entire microbial community using clean raw reads. The sequence similarity (determined using MASH distances) of raw reads revealed clear clustering based on vegetation type (*Cymodocea* vs. *Caulerpa*; Figure 2A), which was consistent with our previous study based on 16S rRNA gene amplicon analysis [17] and validated the use of these data despite low-to-medium coverage. Further, C*aulerpa* metagenomes showed lower distance to centroid (0.04745) compared to *Cymodocea* (0.6494), suggesting that their microbial communities were more homogeneous than those of *Cymodocea*.

**Figure 2.**
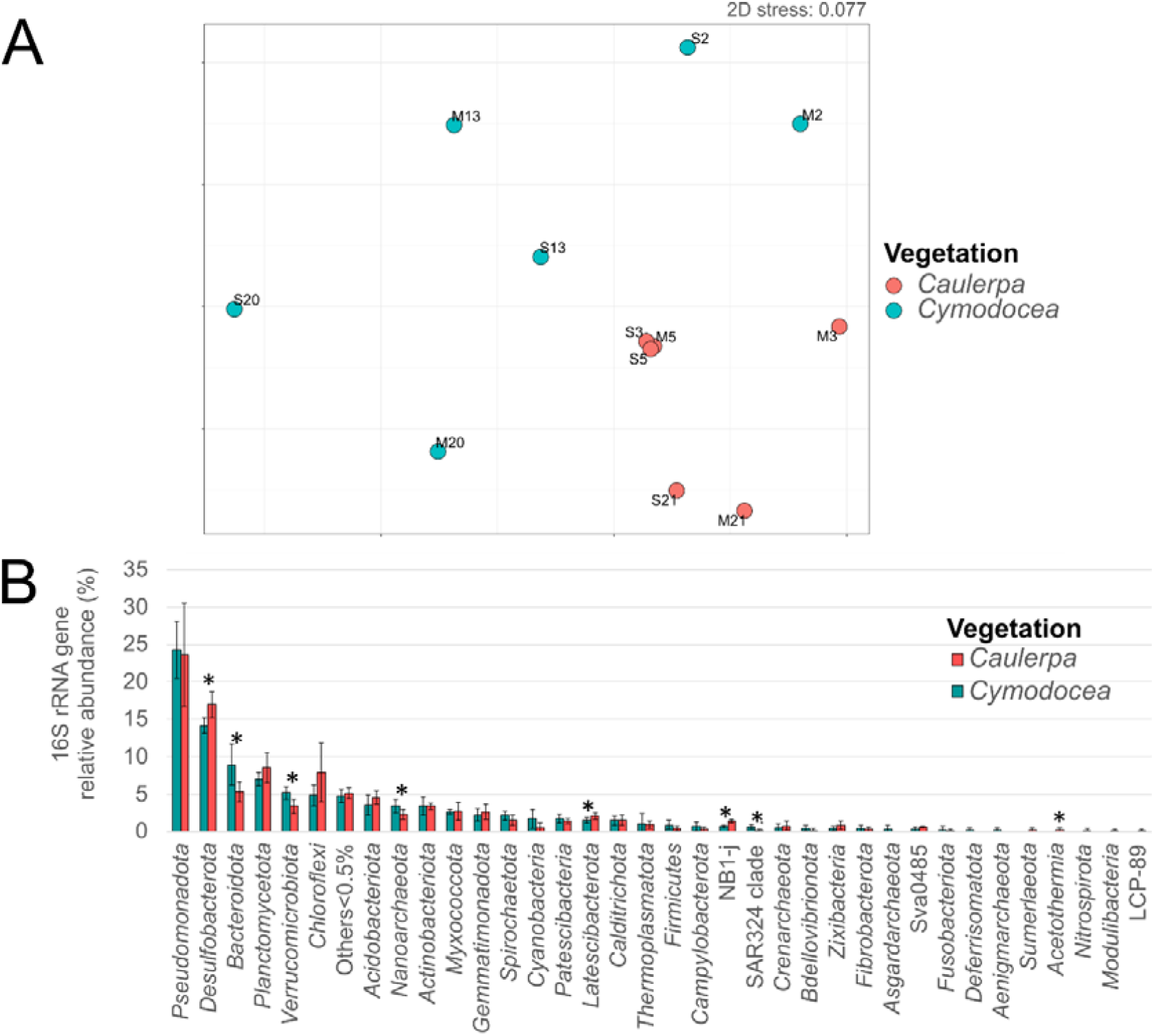
Comparison of microbial communities colonized by *Caulerpa prolifera* (red, n=6) and *Cymodocea nodosa* (blue, n=6). A) Non-metric multidimensional analysis (NMDS) based on MASH distances calculated from metagenomic reads. B) Relative abundance (y-axis) at the phylum level (x-axis) based on 16S rRNA gene reads. Error bars indicate the standard deviation between biological replicates for each vegetation (n=6). Asterisks indicate phyla displaying statistically significant differences based on ANOVA.

Reads corresponding to 16S rRNA genes were extracted from the metagenomic samples and classified against the SILVA database. A total of 101 phyla were detected, with *Pseudomonadota, Desulfobacterota, Bacteroidota*, and *Planctomycetota* being most abundant (Figure 2B). Notably, *Desulfobacterota* was significantly more abundant in *Caulerpa* sediments, whereas *Bacteroidota* and *Verrucomicrobiota* were enriched in *Cymodocea* sediments (ANOVA, p-value < 0.05), in agreement with our previous observations based on 16S rRNA gene metabarcoding [17].

### *Caulerpa*-associated microbiota displays larger potential for sulfate-reduction and carbohydrate metabolism

To examine metabolic differences in detail, KEGG module abundances were quantified using genes predicted from metagenomic contigs. Rarefaction curves using KO-annotated genes (Figure S1) suggested a slightly lower functional diversity in *Caulerpa* samples. This observation aligns with the lower Nonpareil diversity observed for *Caulerpa* compared to *Cymodocea* sediments. In addition, 227 complete KEGG modules (>75% of genes present in each) were identified. Of these, 67 displayed statistically significant differences (ANOVA) between sediment types **(Supplementary Table 1)**, with 65 modules enriched in *Caulerpa* sediments and only two in *Cymodocea* sediments. Enriched modules in *Caulerpa* included pyruvate oxidation, pentose phosphate pathway, TCA cycle, degradation of pyrimidines and aromatic compounds. Sulfate reduction, likely linked to *Desulfobacterota*, was also enriched in *Caulerpa* sediments, supporting the previous hypothesis that microbial species associated with *Caulerpa* are able to stimulate sulfate reduction in the sediments they inhabit [11, 20, 21].

Given that carbohydrates predominate in the biochemical composition of *Caulerpa* and *Cymodocea* [22, 23], and that sediment microbial communities play a key role in degrading decaying organic matter [24], predicted proteins from the metagenome assemblies were analyzed for carbohydrate-active enzymes (CAZymes) as well as proteases and sulfatases. Sediments colonized by *Caulerpa* exhibited a marginally significant higher abundance of CAZymes compared to those with *Cymodocea* (Figure 3A; t-test p-value = 0.069), alongside a slightly higher median CAZyme families diversity per million proteins and lower variance **(Figure 3B)**, although these differences were not statistically significant (Mann-Whitney, p-value 0.4848). Similarly, protease abundance was slightly greater in *Caulerpa* sediments but did not reach statistical significance (t-test p-value 0.8477). Similar abundance of sulfatases was observed between the two sediment types **(Figure 3)**. To evaluate CAZyme compositional differences, exclusive and significantly enriched CAZy families were identified **(Figure S2)**. On one hand, *Caulerpa* had 9 exclusive CAZy families, while *Cymodocea* had 53. On the other hand, a total of 73 CAZy families with statistically significant differences in terms of relative abundance between sediments were identified **(Figure S2)**. The most abundant CAZy families were enriched in *Caulerpa*, while less abundant ones were in *Cymodocea. Caulerpa*-enriched or exclusive CAZy families included CBM40 and GH33 (containing members involved in sialic acid metabolism) and GH50 (containing members with known β-agarase activity). Though agar is typical of red algae, not green algae like *Caulerpa*, we speculate that this GH50 might act on similar polysaccharides, or that GH50 producing strains are able to utilize polysaccharides both from green and red algae. Altogether, these results suggest a higher metabolic potential for organic matter degradation and carbohydrate metabolism for microbial communities associated with *Caulerpa* sediments. These observations are consistent with previous studies showing that *Caulerpa* spp. can substantially modify sediment biogeochemical properties, promoting the accumulation of organic matter and sulfides [11]. Taken together, these findings indicate that the distinct microbial functional profiles are more likely linked to the direct environmental modifications induced by *Caulerpa* than to indirect effects associated with lagoon-wide eutrophication and organic matter enrichment.

**Figure 3.**
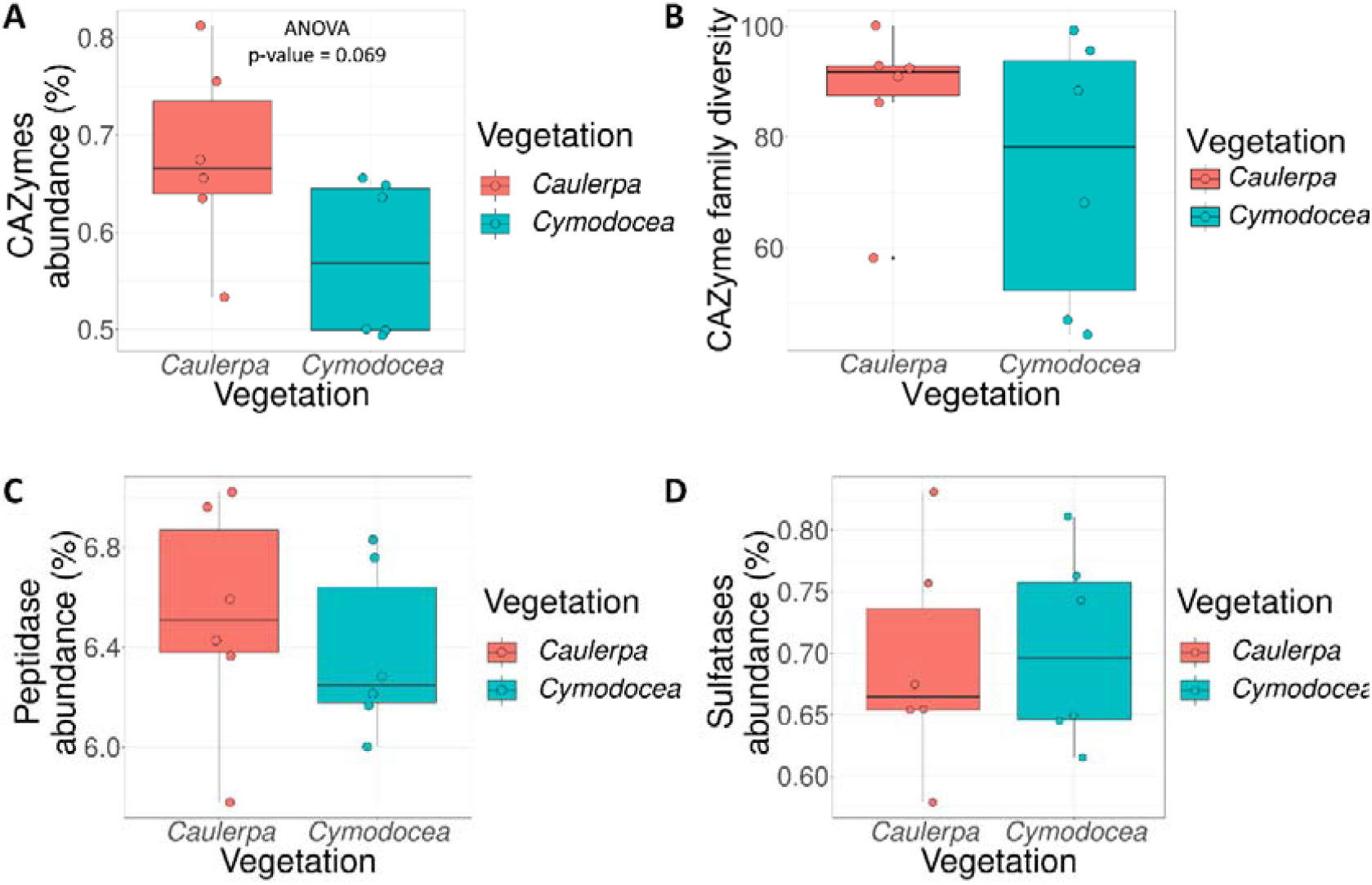
Metabolic potential for organic matter degradation based on the abundance (A) and diversity (B) of predicted CAZymes (measured as number of CAZyme families per million proteins), as well as the abundance of peptidases (C) and sulfatases (D). Boxplots represent the interquartile range (25th–75th percentile), with the median shown as a horizontal line. Whiskers extend to values within 1.5 times the interquartile range, and individual dots represent values from each metagenome (n = 6 per vegetation type; see color legend). Note the slightly higher abundance of CAZymes in *Caulerpa*-associated sediments compared to *Cymodocea*.

### Identification of MAGs involved in the degradation of *Caulerpa* polysaccharides

A total of 158 unique MAGs with completeness above 75% and contamination below 10% (Q50 > 50) were recovered **(Supplementary Table 2)**. These MAGs were primarily classified within *Pseudomonadota, Bacteroidota*, and *Desulfobacterota* phyla **(Figure 4A)**, consistent with earlier phylum-level findings (Figure 2B). Less common phyla like the candidate *Eisenbacteria, Krumholzibacterota, Electryoneota*, and *Blakebacterota* were also recovered as well as four MAGs belonging to the domain *Archaea* (not included in **Figure 4)**. MAGs represented 8.3% to 29.8% of metagenomic reads, with *Caulerpa* metagenomes showing higher percentages *(Figure 4B)*. The higher read recruitment observed in *Caulerpa* metagenomes aligns with the lower alpha diversity in *Caulerpa samples* **(Table 1)**, which favors better assembly and therefore MAG recovery [25].

**Figure 4.**
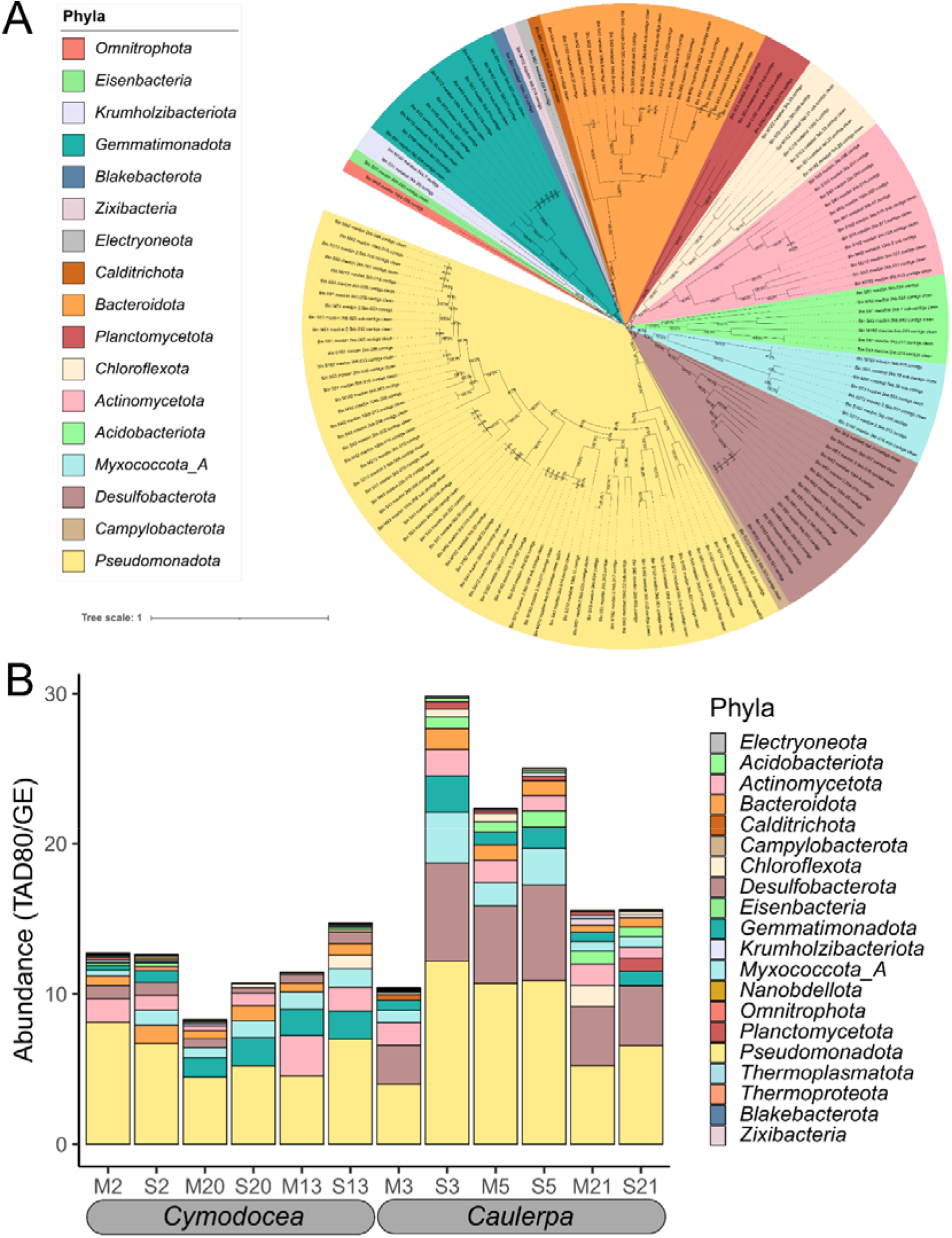
Composition and relative abundance of metagenome-assembled genomes (MAGs) recovered from Mar Menor sediments. (A) Phylogenomic tree of 154 bacterial MAGs constructed from the alignment of 120 universal single-copy marker genes. Taxonomic classification is indicated by branch background color (see legend). (B) Aggregated relative abundance (y-axis), measured as TAD80/GE (see methods), of MAGs grouped by phylum across 12 metagenomes (x-axis). Colors correspond to phyla as shown in panel A. Note the higher overall relative abundance of MAGs in *Caulerpa*-associated sediments compared to *Cymodocea*.

MAGs were categorized as preferentially associated in *Caulerpa, Cymodocea*, or both, based on relative abundance. Criteria included exclusive presence in one sediment type and statistically significant differences in relative abundance (ANOVA). A total of 92 MAGs were common to both, whereas 50 and 16 MAGs were preferentially associated with the *Caulerpa* and *Cymodocea* sediments, respectively (**Figure S3)**. Taxonomically, *Desulfobacterota* MAGs were mainly associated with *Caulerpa*, while *Bacteroidota* were more common in *Cymodocea* **(Figure S3)**, corroborated by abundance data (Figure S3). These results are in agreement with the higher sulfide concentrations found in *Caulerpa* colonized sediments [13] as *Desulfobacterota* are known dissimilatory sulfate reducers.

Differences in CAZyme and protease abundances **(Figure 3)** were also analyzed at the MAG level. MAGs from *Bacteroidota, Planctomycetota* and *Calditrichota* showed >2% of genes dedicated to carbohydrate degradation **(Figure S4**). Absolute CAZyme encoding gene counts also highlighted *Chloroflexota. Archaea* MAGs had the lowest values. For peptidases, *Gemmatimonadota* had the highest relative abundance (∼10%), followed by *Pseudomonadota* and *Bacteroidota* **(Figure S4)**. In absolute terms, *Pseudomonadota, Desulfobacterota*, and *Acidobacteriota* MAGs had the highest number of peptidases. Plotting MAG completeness vs. CAZyme encoding gene counts revealed an enrichment of these genes in *Bacteroidota, Planctomycetota*, and *Chloroflexota* MAGs **(Figure S5A**). Most of them belonged to the *Caulerpa* group **(Figure S5B)**. This trend held whether analyzing absolute CAZyme encoding gene counts or relative to total genes.

Furthermore, a heatmap of CAZyme encoding gene abundance per MAG showed a cluster of glycosyl hydrolases enriched in these *Bacteroidota* and *Planctomycetota* MAGs associated with *Caulerpa* **(Figure 5A)**. Such glycosyl hydrolases might act on pectin **(Figure 5B)** and we hypothesize that they are likely involved in the degradation of pectin-like molecules present in the Mar Menor sediments. Pectin is a complex plant polysaccharide composed of homogalacturonan, rhamnogalacturonan I & II, and xylogalacturonan [26]. While pectin has not been found in *Caulerpa*, similar sulfated polysaccharides have been reported [27–29]. Therefore, these enzymes likely target structurally analogous compounds. Despite the functional potential for pectin-like degradation seems to exist for MAGs associated with *Caulerpa* sediments, this does not preclude the presence of MAGs with similar capabilities in *Cymodocea* sediments but not recovered in our dataset.

**Figure 5.**
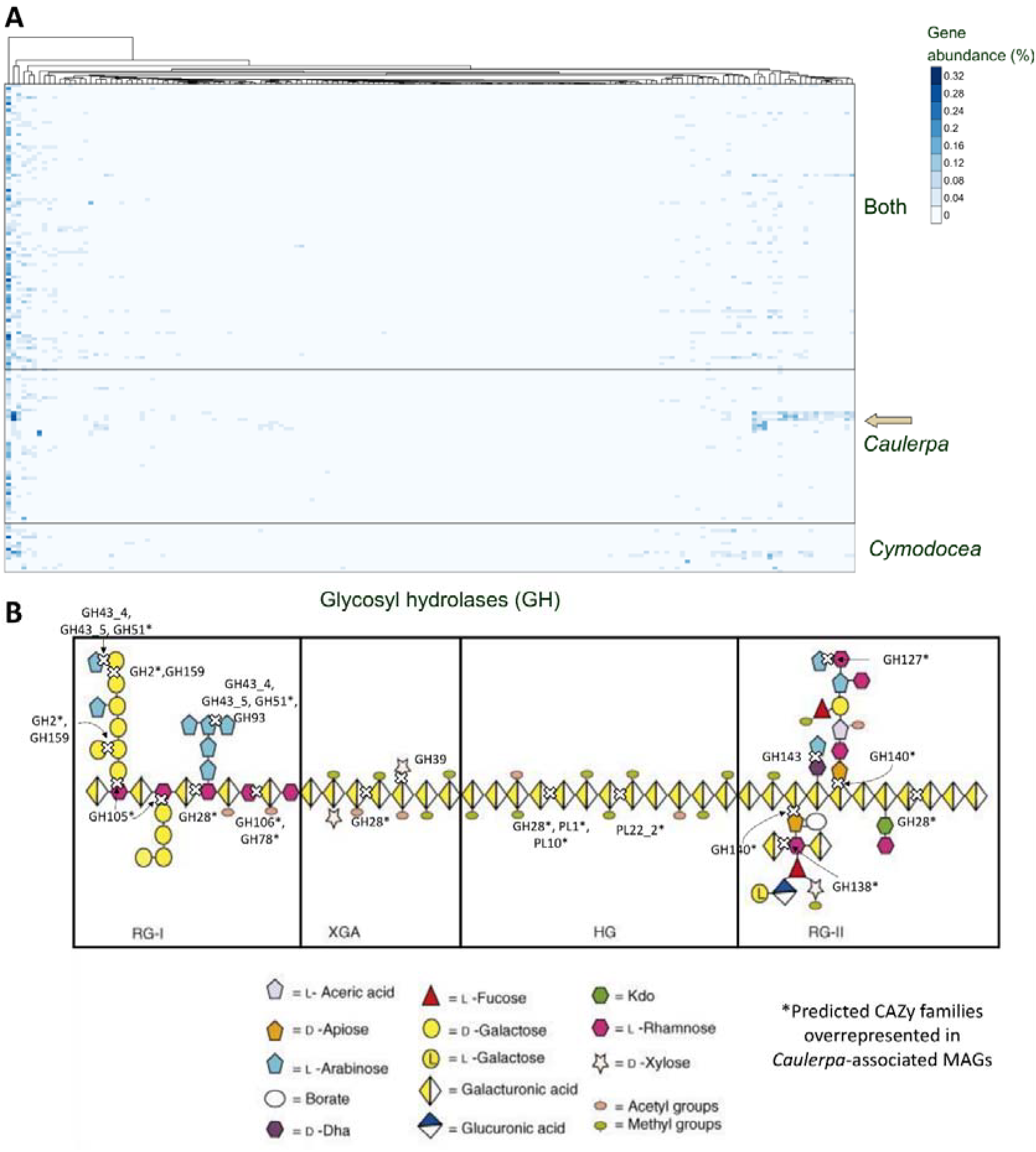
Metabolic potential for carbohydrate modification encoded by metagenome-assembled genomes (MAGs). (A) Heatmap showing the proportion of putative proteins annotated with glycosyl hydrolases families (x-axis) for each MAG (y-axis), with color intensity (blue scale) indicating increasing percentages of gene abundance. MAGs are grouped by their association with *Caulerpa, Cymodocea*, or both sediment types. *Caulerpa*-associated MAGs enriched in GH are marked with a green arrow. (B) Diagram of pectin and its four main subunits—Rhamnogalacturonan I (RG-I), Xylogalacturonan (XGA), Homogalacturonan (HG), and Rhamnogalacturonan II (RG-II). Predicted CAZy families and subfamilies involved in pectin degradation are shown, with those enriched in *Caulerpa-*associated MAGs (as identified in panel A) indicated by asterisks. Note the broad distribution of enriched CAZymes across the pectin structure. Panel B is adapted from Mohnen 2008 [26].

### The most abundant MAGs were not the most actively transcribed

To assess transcriptional activity, metatranscriptomic reads were mapped to the MAGs predicted genes, and expression was normalized by gene length and transcriptome size. Plotting MAG abundance versus aggregated gene expression for each MAG revealed that the most abundant MAGs were not necessarily the most transcriptionally active **(Figure 6)**, revealing a lack of correlation between relative abundance and transcriptional activity that was statistically corroborated by a low Spearman’s rho coefficient (ρ=0.067). For instance, *Desulfobacterota* MAGs dominated in relative abundance but were not the most transcriptionally active members of the community. Instead, certain *Bacteroidota, Pseudomonadota*, and *Campylobacterota* MAGs exhibited high aggregated gene expression, mainly driven for certain genes including elevated transfer-messenger RNA (tmRNA) levels, which are involved in resolving stalled ribosomes and targeting faulty proteins for degradation [30]. Indeed, in Figure 6, within the 59 MAGs/site with top expression levels (95% percentile), only 7 rank in the 95% percentile of relative abundance. This proportion increases when lower, albeit still high, levels of transcription are considered. The lack of correlation between the abundance in metagenomes and activity in metatranscriptomes was also observed at the gene level. Metagenomic studies typically focus on the most abundant MAGs under the assumption that these are the key drivers of the community and low abundant MAGs have a marginal impact on ecosystem functioning. Our results, however, challenge such assumption and emphasize the need of activity-based data (e.g., metatranscriptomics, metaproteomics) for a more accurate assessment of the relative importance of each species in ecosystem functioning.

**Figure 6.**
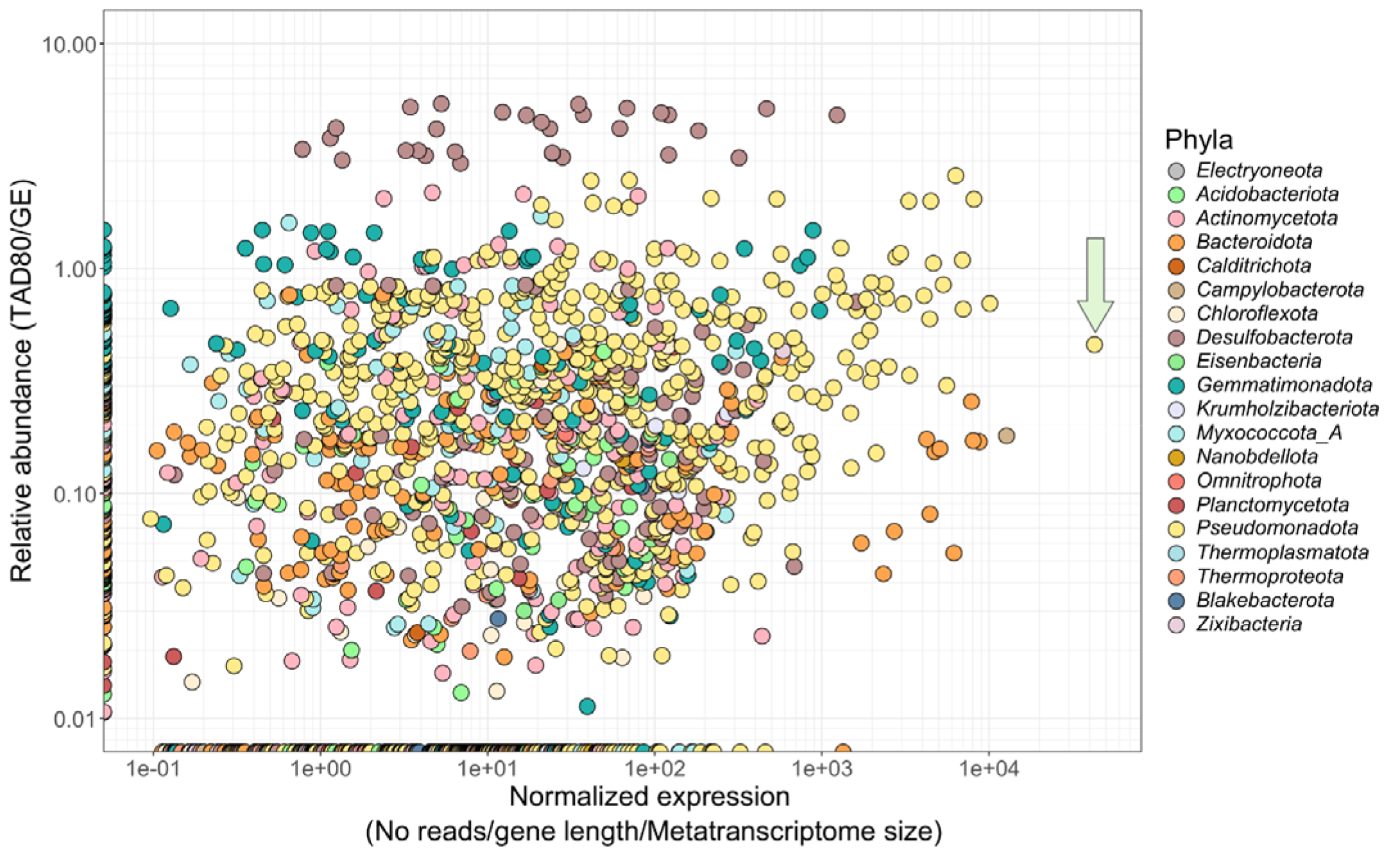
Relationship between transcriptional activity and relative abundance of metagenome-assembled genomes (MAGs). Dot plot showing MAG relative abundance in the metagenome (y-axis, log scale) versus their aggregated normalized gene expression in the corresponding metatranscriptome (x-axis, log scale). Each dot represents a single MAG in a specific sediment sample. Colors indicate the MAG’s taxonomic affiliation at the phylum level (see legend). The arrow vertical highlights the MAG with the highest overall gene expression in the dataset. Note that the most abundant MAGs, such as those from *Desulfobacterota*, do not exhibit the highest transcriptional activity.

### Most highly expressed genes in MAGs correspond to unannotated proteins

Among the 38,743 expressed genes, 30.7% corresponded to unannotated proteins. When the 100 most expressed genes were considered, the proportion of unannotated proteins dramatically increased to 97%. Thus, the function of the most expressed proteins in these sediments is basically unknown.

At the MAG level, one *Gammaproteobacteria* MAG (Bin_S43_maxbin_2kb.011.contigs) showed the highest expression **(Figure 6, arrow)** mainly due to a single unannotated gene. This gene, encoding a protein that we named very highly transcribed gene product (hereinafter VHTgp), was present in all metagenomes (>98.6% identity) and expressed in two metatranscriptomes (S20 and M2, corresponding to *Cymodocea* colonized sediments) at very high levels compared to all other genes (it was the most highly expressed gene in S20 and the sixth most highly expressed in M2). It was located between the mlaABCDEF operon (a set of genes that are co-transcribed together and are involved in lipid membrane asymmetry [31]) and a DUF302 gene encoding a conserved protein family of unknown function found in both *Archaea* and *Bacteria* **(Figure S6)**. The uncharacterized gene was transcribed in the same direction as that encoding DUF302 but opposite to the *mlaABCDEF* operon, suggesting that the gene may be co-transcribed with that of DUF302 but not with the *mlaABCDEF* operon. Similarity searches (i.e., BLASTp against NCBI NR) revealed the existence of contigs with similar genetic structure in other *Gammaproteobacteria* MAGs recovered from coastal marine sediments in China and Australia, as well as from a freshwater lake in northern United States **(Figure S6)**, indicating its widespread nature. In addition, conserved domain analysis pointed to distant similarities to conserved domains in periplasmic binding proteins and glycosyltransferases. Structural predictions based on amino acid sequences were used to search for structural matches and infer possible functions, ligands, and cellular locations. The top 5 PDB hits were all very distantly related to glycosyl hydrolases, sharing between 8 – 25% sequence identity with VHTgp: GH14 (1B1Y), GH106 (5MQM), GH35 (7VKW), GH13 (1QHO), GH42 (4UNI) with diverse activities such as β-amylase, α-L-rhamnosidase, β-1,2-glucan synthase, α-amylase, and β-1,6/β-1,3-galactosidase (TM-scores = 0.47 – 0.50). Therefore, polysaccharide and ion binding were predicted as possible ligands with a low confidence score of 47% (C-score). Hence, we hypothesized it could be a periplasmic protein capable of polysaccharide modification.

### Recombinant expression of VHTgp reveals peptidoglycan degradation activity

Structural models and other topology predictors postulate VHTgp to have a poor 3-dimensional fold. The ESM fold model predicted with low confidence (pLDDT 20 - 40% across the entire protein) a few α-helices (10%) and β-sheets (17%) but the global structure was poorly identified (30% with no predicted secondary structure) **(Figure S7A-C)** [32, 33]. Additionally, a transmembrane helix (122 – 144) was identified by MEMSAT-SVM, consequently predicting residues 145 – 223 (C-terminal) as extracellular **(Figure S7D)**. This was also supported from the COFACTOR results, as the protein was predicted with high confidence to be in the extracellular region. Collectively, these results suggested VHTGP may be either unstable in vivo or bound to the membrane hinting to possible difficulties with expression.

Recombinant VHTgp was difficult to express and purify in a heterologous system. Nevertheless, SDS–PAGE analysis of E. coli cultures after overnight induction revealed a strong overexpression band at approximately 25 kDa, below the predicted molecular weight of 27 kDa **(Figure S8)**, suggesting C-terminal truncation of the protein. After clarification of the cell lysate, both soluble and insoluble fractions contained the over-expressed protein which is in-line with bioinformatic predictions as a possible membrane-bound protein. Immobilized metal affinity chromatography (IMAC) purification of the clarified cell lysate failed perhaps due to the pre-mature cleavage of the (His)_6_-tag C-terminal of the protein **(Figure S9)**. Optimization yielded no significant improvements in expression (outlined in Materials & Methods).

To further investigate the hypothesis that the enzyme functions as a polysaccharide-modifying protein potentially associated with the cell membrane, we tested whether it exhibits peptidoglycan-degrading activity. This was assessed using a turbidity reduction assay (TRA) with permeabilized E. coli cells [34]. Clarified cell lysates showed a decrease in optical density (OD_600nm_), indicative of peptidoglycan degradation activity **(Figure 7A, red)**. This suggests that the VHTgp protein found in the cell lysate degrades the peptidoglycan either through the polysaccharide as a CAZyme (hydrolytic lysozyme or lytic transglycosylase [35]) or the peptide crosslinker as an amidase or endopeptidase. Additionally, after boiling the cell lysate we observed a disappearance of activity **(Figure S10)**. To determine the type of enzyme (CAZyme vs amidase/endopeptidase) we subjected it to a secondary assay that relies on the fluorescence liberated from the oligosaccharide cleavage of peptidoglycan conjugated with fluorescein. Clarified cell lysates were tested for activity, yet no fluorescence was observed indicating VHTgp is most likely not a CAZyme but degrades the peptidoglycan as an amidase or endopeptidase via the peptide crosslink, contrary to previously discussed predictions **(Figure 7B)**.

**Figure 7.**
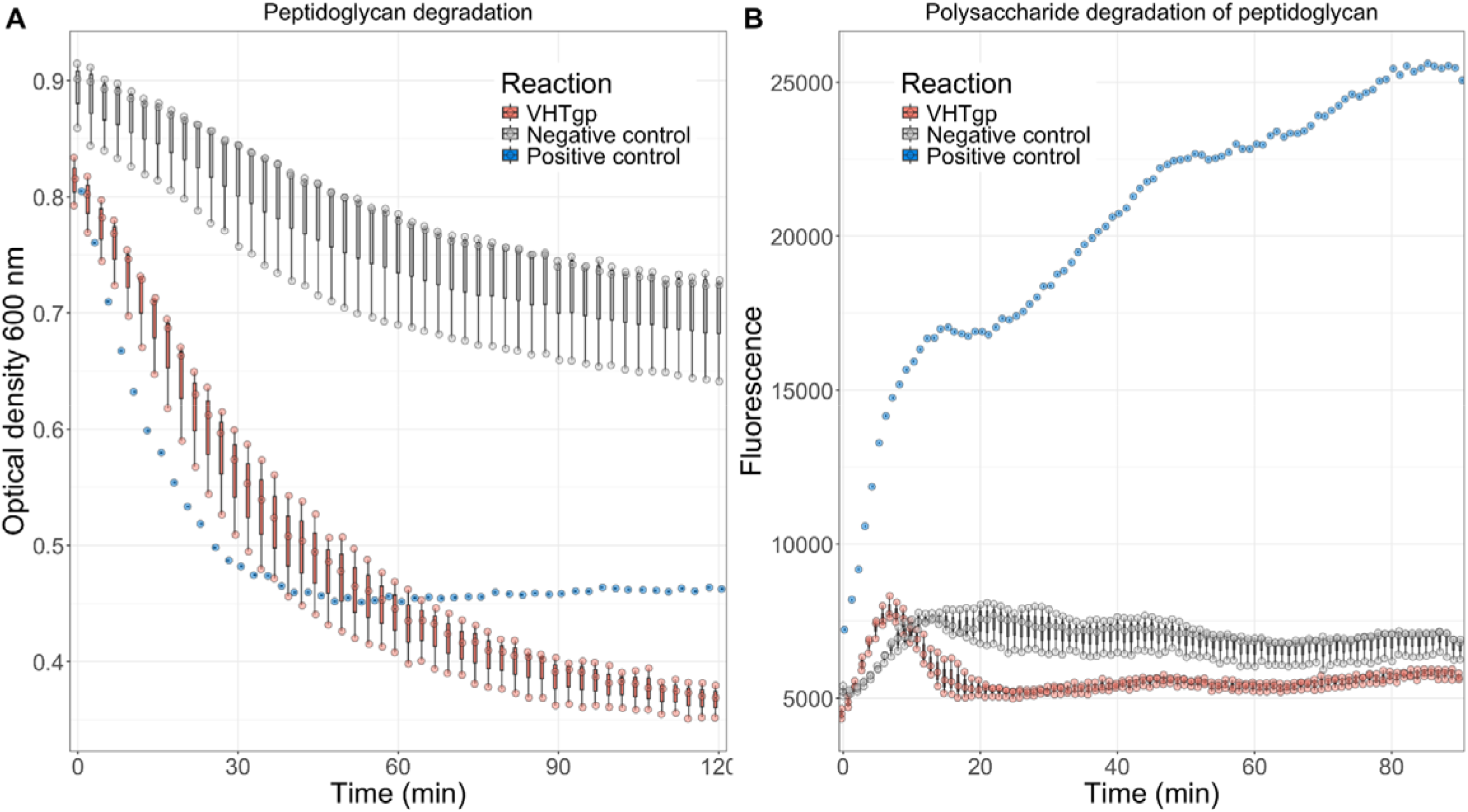
A) Turbidity Reduction Assay highlights general peptidoglycan degradation. Clarified cell lysates containing expressed VHTgp (red) protein show positive peptidoglycan degradation activity of permeabilized *E. coli* cells while control clarified cell lysates containing an empty pET28a plasmid (negative control, grey) remains inactive. B) EnzChek Lysozyme Assay reveals polysaccharide degradation of the peptidoglycan. Both clarified cell lysates containing expressed VHTgp (red) protein and empty pET28a plasmid (negative control, grey) show no polysaccharide degradation activity of fluorescein conjugates to peptidoglycan. In both experiments, HEWL (lysozyme that degrades the polysaccharide of the peptidoglycan) is used as a positive control (blue). Each dot shows the value for each of the three biological replicates per reaction.

These types of proteins belong to a class of bacterial autolysins responsible for peptidoglycan turnover during cell growth and division. Therefore, the protein encoded by the VHTgp gene is likely involved in remodeling endogenous peptidoglycan within the cell wall rather than in the degradation of exogenous peptidoglycan released from dead bacterial cells. Cell wall amidases include N-acetylmuramoyl-L-alanine amidases, that cleave the peptide stem directly from the oligosaccharide while peptidases are endo- and/or carboxypeptidases [36]. The upregulation of the gene encoding this protein may be due to the proliferation of this species at the time of sampling due to an abundance of available carbon source and in turn leading to a rapid onset of growth and division. Alternatively, it could be involved in damage-induced cell-wall reparation given that stress sigma factors, such as *rpoH* or *rpoS* encoded by this MAG, were also expressed at the same time than VHTgp.

### Concluding remarks

Our study reveals a clear predominance of sulfate-reducing bacteria and a higher potential for organic matter degradation in sediments colonized by the green macroalgae *Caulerpa* compared to those associated with the seagrass *Cymodocea*, thus confirming our initial hypothesis. Notably, we identified several MAGs with the potential to breakdown the pectin-like polysaccharide characteristic of the *Caulerpa* cell wall. Mechanistically, we propose that the higher organic matter content of *Caulerpa* sediments [16] promotes oxygen depletion at the sediment surface by aerobic bacteria including polysaccharide-degraders, thereby favoring the proliferation of anaerobic taxa, particularly sulfate-reducing bacteria. This leads to increased sulfide concentrations, which are toxic for *Cymodocea*. Under this framework, the reported negative impact of *Caulerpa* on *Cymodocea* [10, 15], if present [16], would be indirectly mediated by shifts in the sediment-associated microbial community rather than by direct plant–plant interactions.

Our metatranscriptomics data indicate that less abundant microbes should not be discounted regarding biogeochemical cycling since the metagenomically dominant microbes identified in our study were not the most active in terms of gene transcription. We would like to argue that temporally resolved sampling would be the best way to investigate the discrepancies of metagenomic abundance and metatranscriptomics activities.

Finally, we provide experimental support for the functional annotation of a widely distributed, highly expressed hypothetical gene, encoding a novel peptidoglycan degrading enzyme, likely involved in cell wall remodelling. This gene, associated with a *Gammaproteobacteria* MAG, was detected across multiple sites in the Mar Menor lagoon at both sampling times (March and September), indicating broad spatial distribution and temporal stability. The absence of close homologs in public databases highlights its novelty. We anticipate that the heterologous expression and functional characterization presented here, together with its deposition in public databases, will facilitate future investigations of this enzyme. More broadly, our results emphasize the strong need to study at least the highly expressed fraction of the many hypothetical genes found in metagenomic datasets. We would like to advocate an integrative approach that combines multi-omics with targeted protein expression and functional characterization, as this strategy would advance the knowledge of the functional potential and ecological role of environmental microorganisms.

## Material and methods

### Study area

Mar Menor is a shallow marine coastal lagoon (maximum depth 7m, average depth 4.5m) located in the Region of Murcia, in the southeast of Spain. It is separated from the Mediterranean Sea by a narrow barrier of land called “La Manga” although there are several inlets through which water exchanges between them. By this reason, water volume is relatively confined with a renewal time of about 1 year. Salinity and temperature range from 4.2% to 4.7% and 10 to 31°C, respectively. In the last decades, the lagoon has experienced drastic and sudden changes due to the pressure exerted by several anthropic factors such as hydrological changes, mining, agriculture activities and contaminants of emerging concern (pharmaceuticals, pesticides, etc.) [37].

### Sampling design

We collected sediment samples from 6 different stations distributed along the Mar Menor, 3 of them with presence of *Ca. prolifera* (P3, P5 and P21) and 3 with *Cy. nodosa* (P2, P13, and P20) **(Figure 1)**. Samples were collected at two different times, March and September 2018. Thus, the dataset is composed of 6 samples for each vegetal species. Sediment samples were collected from the upper 4.5 cm, weighted, homogenized with 50 mL of PBS 1 × 3.5% NaCl and frozen at -80°C until processing. It should be noted that these samples were previously described and analyzed using 16S rRNA gene amplicon sequencing in [17], except the stations P20 and P21 that were not included in the previous paper.

### DNA/RNA extraction and sequencing

DNA and RNA were simultaneously extracted from 2 grams of unfrozen sediment with the RNeasy Power Soil Total RNA Kit (Qiagen) and its complementary RNeasy Power Soil DNA Elution Kit (Qiagen) following manufacturer instructions. Extracted nucleic acids concentrations were measured using the Qubit dsDNA High Sensitivity (HS) and RNA HS kits (Invitrogen). RNA extractions were treated with DNase for 1 hour at 37ºC and absence of DNA was confirmed with PCR using 16S rRNA gene universal primers. Sequencing was performed on an Illumina NovaSeq6000 run (2 × 150 bp) in the National Centre for Genomic Analyses (CNAG-CRG, Barcelona).

### Raw reads processing

Raw reads were trimmed and adaptors removed using Trimmomatic v0.36 (LEADING:3 TRAILING:3 SLIDINGWINDOW:4:15 MINLEN:36) [38]. Read quality was then checked with FastQC v0.11.9 [39]. To estimate the sample coverage and diversity of cleaned reads Nonpareil v3.303 was employed (-T kmer -f fastq -X 50000) [40]. Distances between raw reads calculated with MASH v1.1 [41] were used to draw a PCA plot in R studio [42, 43] using the libraries “vegan” and “factoextra” [44, 45]. rRNA gene reads were extracted using SortMeRNA v4.2.0 [46] and 16S rRNA gene reads were classified taxonomically by BLASTn v2.9.0+ against SILVA database r138.1 [47] (coverage>70%, identity>83%, threshold previously proposed for Classes [48]). Regarding read assembly, a preliminary test comparing MEGAHIT v1.2.9 [49] and IDBA-UD [50] was carried out and MEGAHIT was chosen given the larger number of contigs and contig lengths.

### Assembly analyses

Proteins were predicted from contigs using Prodigal v2.6.3 and the option “-p meta” [51]. Carbohydrate-active enzymes (CAZy) were annotated using DIAMOND search for BLASTp hits in the CAZy database (http://www.cazy.org/)[52] (coverage > 50%, identity > 40%) and HMMER search against the dbCAN CAZy domain HMM database (e-value < 1e-15, coverage > 0.35) [53, 54]. Only proteins annotated by both tools were considered, as previously suggested [55]. Peptidases and sulfatases were searched in MEROPS [56] and SulfAtlas databases [57], respectively, using BLASTp (coverage > 50%, identity > 40%). Anvi’o-7.1 [58] was employed to annotate the assembly against the KEGG KO database [59, 60] and calculate the number of KO per KEGG module and module completeness using the “anvi-estimate metabolism” program. Statistical differences were tested by ANOVA using the aov function of the package “vegan” in R.

### MAG analyses

Binning recovery was maximized for each metagenome assembly employing two different tools, MaxBin2 v2.2.7 [61] and MetaBAT2 v2.15 [62], and different minimum contig size [1kb (only MetaBAT2), 1.5 kb (only MaxBin2), 2kb, 2.5kb, 3kb, 5kb and 10 kb]. The optimum size for each tool on each metagenome was selected for further analyses. Based on checkM results [63], the criteria to select the optimum contig size were i) highest number of bins with completeness > 80% and contamination < 5%, if equal ii) highest number of bins with completeness > 80% and contamination < 10%, if equal iii) highest number of bins with completeness > 80% and, if still equal iv) the largest contig size was selected. Selected MAGs were then processed through DAS Tool v1.1.3 [64]. Additionally, to filter out low-quality MAGs, completeness and contamination was estimated by checkM v1.1.3 (lineage_wf) and only those with %[completion] ⍰−⍰5*[%contamination]⍰>⍰ 50 were retained for downstream analyses. Highest-quality MAGs (completeness > 80%) were manually refined by checking taxonomy of predicted ORFs (BLASTp against NR database) and sequencing depth (coverage>70%, identity>95%), as previously suggested [65]. Finally, MAGs were dereplicated at 99% ANI using dRep v3.4.2 [66] which kept 159 MAGs.

MAG taxonomy was assigned by GTDB-tk v1.6.0 release 202 [41, 51, 53, 67–71]. The alignment of 120 single-copy marker genes generated by GTDB-tk was used to calculate a phylogenomic tree with FastTree v2.1.10 (options -gamma -lg) [71], which was then visualized in the interactive Tree of Life (iToL) [72].

To estimate MAG abundances in each metagenome, the 80% central truncated average sequencing depth (TAD80)/genome equivalent (GE) ratio was calculated, as previously described [73]. Briefly, reads were aligned against each MAG using bbmap (idtag=t minid=0.95 idfilter=0.97) [74], samtools v1.7 converted and sorted the SAM format file into a BAM format file [75], then the genomecov workflow (-bga) of the Bedtools package v2.30.0 [76] calculated the sequencing depth for each base which was finally filtered by the “BedGraph.tad.rb” (option: -r 0.8) script of the enveomics collection [77] to obtain the 80% central truncated average sequencing depth (TAD80). The number of genome equivalents offers an estimate of the number of prokaryotic cells sequenced in the metagenome based on the abundance of an essential single-copy genes set and was calculated for each metagenome using MicrobeCensus v1.1.1. [78].

Based on the abundance data, three different groups of MAGs were identified: MAGs present in both type of sediments, MAGs associated with *Caulerpa* and *Cymodocea* sediments, respectively. These groups were defined based on the presence/absence of MAGs but also on statistically significant different abundances reported by ANOVA tests.

Predicted proteins from MAGs were functionally annotated with Prokka [79] and interproscan [80] using the Pfam, CDD, SMART and TIGRFAM databases [81–84]. CAZy and peptidases were also searched as described above.

### Metatranscriptomic analyses

Metatranscriptomes were cleaned using Trimmomatic, as described above for metagenomes. Then, rRNA reads were identified with SortMeRNA [85] and extracted to generate rRNA-free metatranscriptomes. To calculate the genes relative expression, rRNA-free metatranscriptomes were aligned against all predicted genes from MAGs using parallel [86] and BLASTn (filtered by best hit, coverage>70%, identity>97%) [87]. The number of mapped transcripts per gene was calculated with the “BlastTab.sumPerHit.pl” script of the enveomics collection and the breadth coverage of each gene was calculated with a homemade Linux script. To filter out genes that might be only mapped on conserved domains, only those genes with breadth coverage equal or higher than 90% were considered as expressed. The normalized expression of each gene was calculated as follows: number of transcripts/gene length/metatranscriptome size.

### Enzyme production

The gene encoding VHTgp was synthesized by Biomatik Limited (Cambridge, ON, Canada) with codon optimization for the expression in *E. coli* and cloned in a pET23a(+) containing a C-terminal (His)_6_-tag and pET-28a(+) vectors containing a N-terminal (His)_6_-tag, thrombin cleavage site and a T7 tag leader peptide. Additionally, an empty control pET28a(+) plasmid was transformed and expressed alongside the recombinant plasmid. E. coli C41(DE3) cells were transformed with both plasmids separately and were cultured overnight (20°C, 180 rpm) in 50 mL auto-inducible medium ZYM5052 supplemented with 50 μg/mL kanamycin or 100 μg/mL ampicillin. We observed no overexpression band when VHTgp was cloned in pET28a **(Figure S8, S9)**. The cells of were harvested by centrifugation (30 minutes at 10,000 rpm, 10°C) and resuspended in Tris/HCl (20 mM, pH7), NaCl (200 mM) buffer to a final concentration of OD_600nm_=80. Both cells were lysed by sonication (30% amplitude, 5 cycles of 20 s ON, 60 s OFF) using a 3 mm tip and the crude cell lysates were collected after centrifugation (30 minutes at 10,000 rpm, 10°C). The cell lysates were kept at 4°C for only a few hours before screening for activity.

Troubleshooting of the expression included a co-expression with pGroEL/ES in ZYM5052 supplemented with 32 μg/mL of chloramphenicol and 0.05% L-arabinose along with appropriate antibiotics of VHTgp. Additionally, a final attempt of optimization included the addition of IPTG (0.5 mM) in LB medium and was agitated overnight (20°C, 180 rpm).

### Activity screening

The peptidoglycan degradation activity of the protein in the clarified cell lysate was quantified by monitoring the OD_600nm_ reading every 2 minutes for 120 minutes at 30°C with a TECAN Infinite 200Pro microplate reader while shaking linearly for 4 s at a 3 mm amplitude before each read in a Greiner 96 Flat Bottom Transparent Polystyrene microplate. The outer membrane of *E. coli* XL1-Blue cells was permeabilized by CHCl_3_-saturated PBS-buffer as previously described^1^ and Tris/HCl (20 mM, pH7) was the final buffer the permeabilized cells were resuspended in. The reaction was initiated when 100 μL of permeabilized cells were added to 100 μL of clarified cell lysate (OD_600nm_=80) to make the clarified cell lysate concentration OD_600nm_=40 and the final buffer composition was Tris/HCl (20 mM, pH7), NaCl (100 mM). Both VHTgp and control empty pET28a clarified cell lysates were tested in biological triplicates. As positive control, we used a stock solution of HEWL diluted to a final concentration of 0.01 mg/mL in the clarified cell lysate containing the empty pET28a plasmid.

The Lysozyme EnzChek Assay (Invitrogen, Thermofisher Scientific) was used to probe carbohydrate degradation activity from peptidoglycan substrate. A stock solution (50 μg/mL) of the provided kit substrate, fluorescein conjugated to M. lysodeikticus peptidoglycan, was prepared in a Tris/HCl (20 mM, pH7) buffer and 50 μL was added to 50 μL of clarified cell lysate to make the final concentration of cell lysate OD_600nm_=40 and final buffer composition as Tris/HCl (20 mM, pH7), NaCl (100 mM). The fluorescence (λ_ex_=485 nm, λ_em_=530 nm) was monitored every 90 seconds over 90 minutes at 30°C using a TECAN Infinite 200Pro microplate reader using Thermo Fisher Scientific Black Nunc 96 Flat bottom polystyrene plates. Both VHTgp and control empty pET28a crude cell lysates were tested in biological triplicates.

## Data availability

Metagenomes and MAGs generated during this study have been deposited in NCBI under BioProject <u>PRJNA901668</u>. Metatranscriptomes are available in NCBI under BioProject PRJNA1397545.

## Acknowledgements/funding

This research was funded by European Union’s Horizon 2020 framework program BLUETOOLS grant (No. 101081957). BA-R was supported by an ACIF18 fellowship (ACIF/2018/179) for his doctoral thesis and received funding through the BEFPI2021 program (BEFPI/2021/004) to undertake a research stay at the Max Planck Institute for Marine Microbiology (Bremen), both funding schemes were provided by the Generalitat Valenciana.

