## Supplementary Material for "Anthropogenically induced shifts in sediment vegetation impact coastal benthic microbiomes"

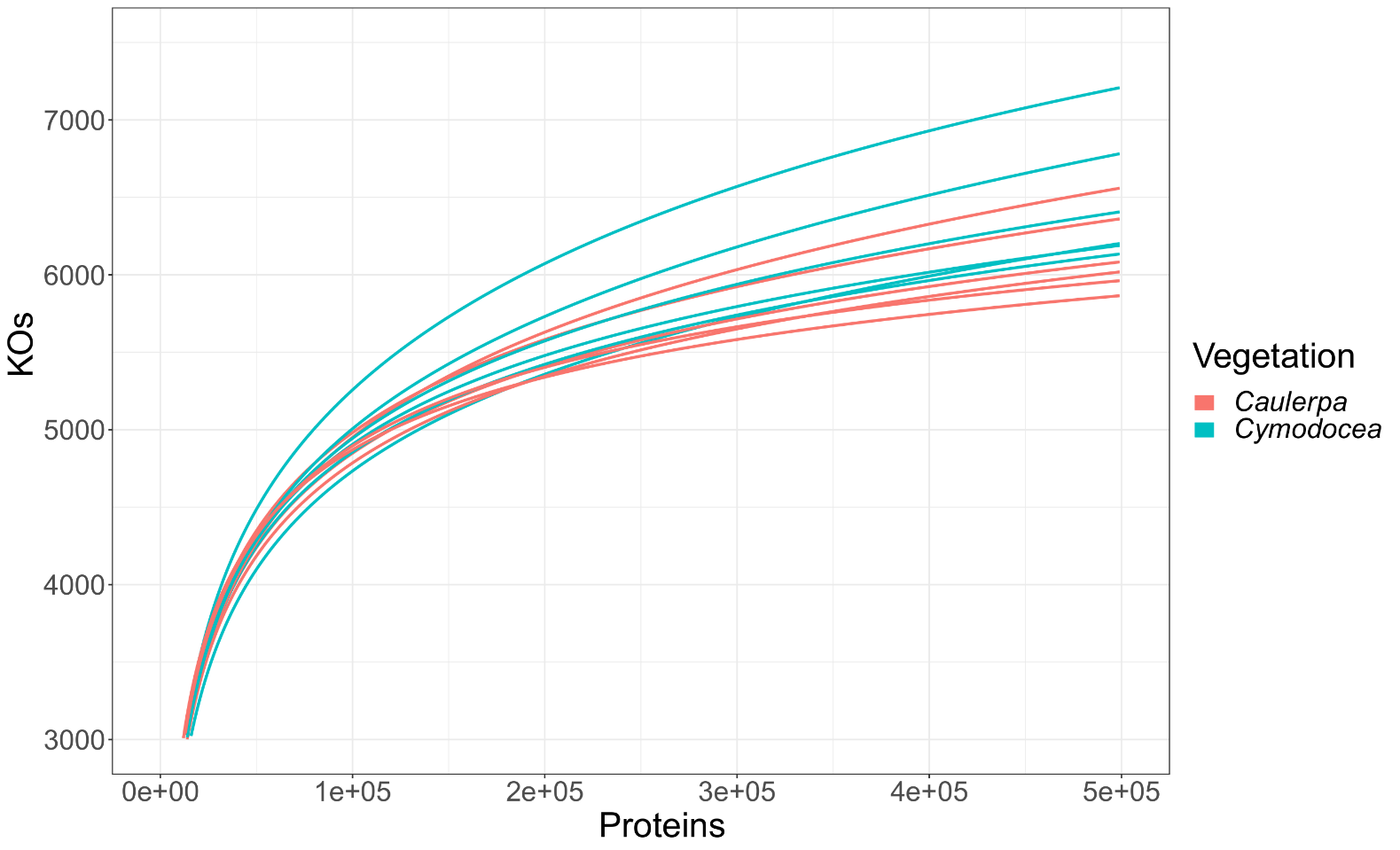

**Figure S1**. Rarefaction curve showing the number of annotated genes with Kegg Orthology numbers (y-axis) respect to the number of proteins (x-axis) for each assembled metagenome (lines). Colors indicate the vegetation covering the sediment, that is, *Caulerpa prolifera* or *Cymodocea nodosa* (see legend).

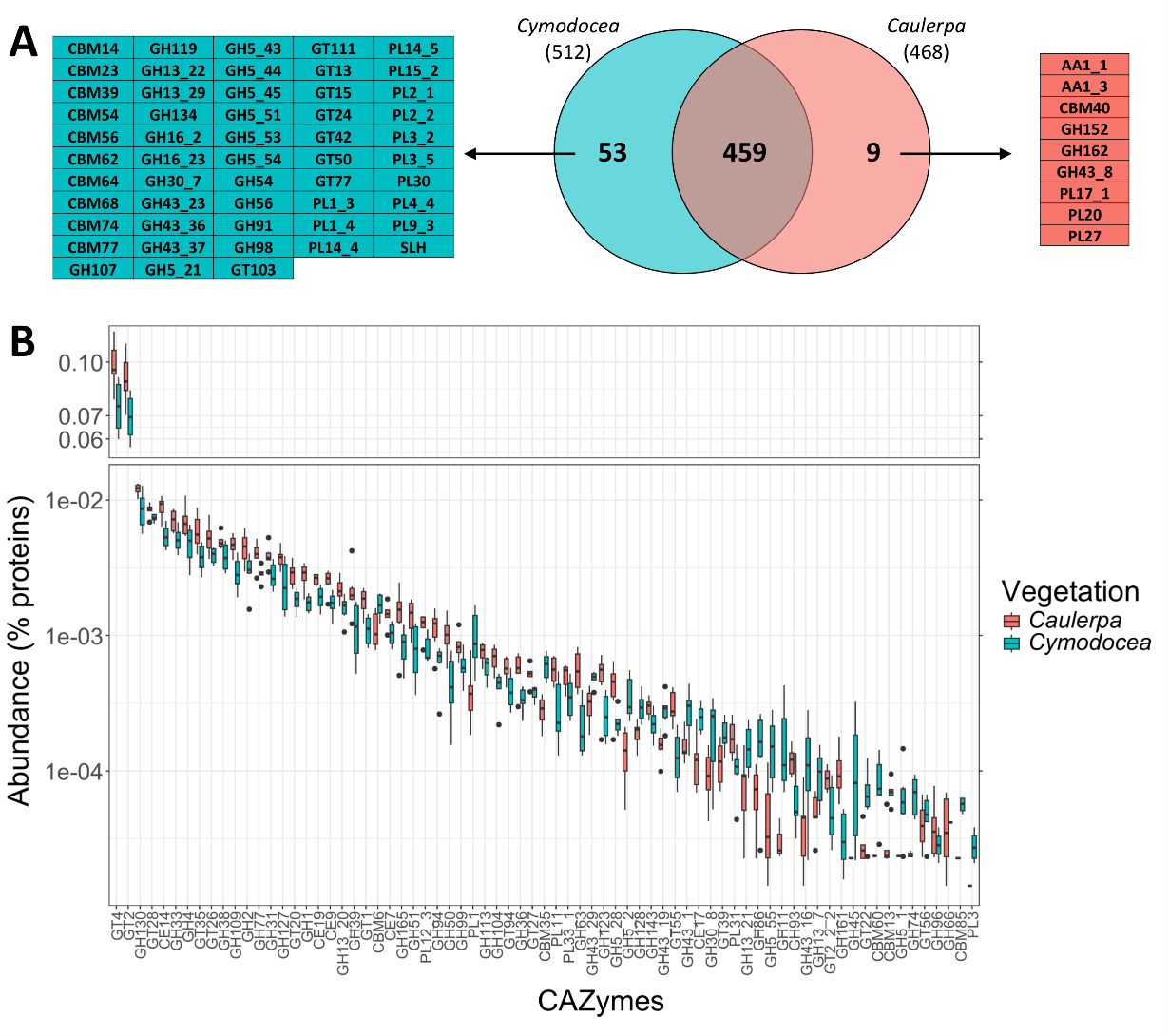

**Figure S2.** CAZyme compositional differences between *Caulerpa* and *Cymodocea* metagenomes.A) Venn diagram showing the number of shared and vegetation-specific CAZy families. For the latter, the CAZyme families and subfamilies are shown next to the Venn diagram, B) Boxplot showing the relative abundance, calculated as percentage of proteins (y-axis), of CAZyme families and subfamilies (x-axis) showing statistically significant enrichment in either *Caulerpa* or *Cymodocea* metagenomes (colors)

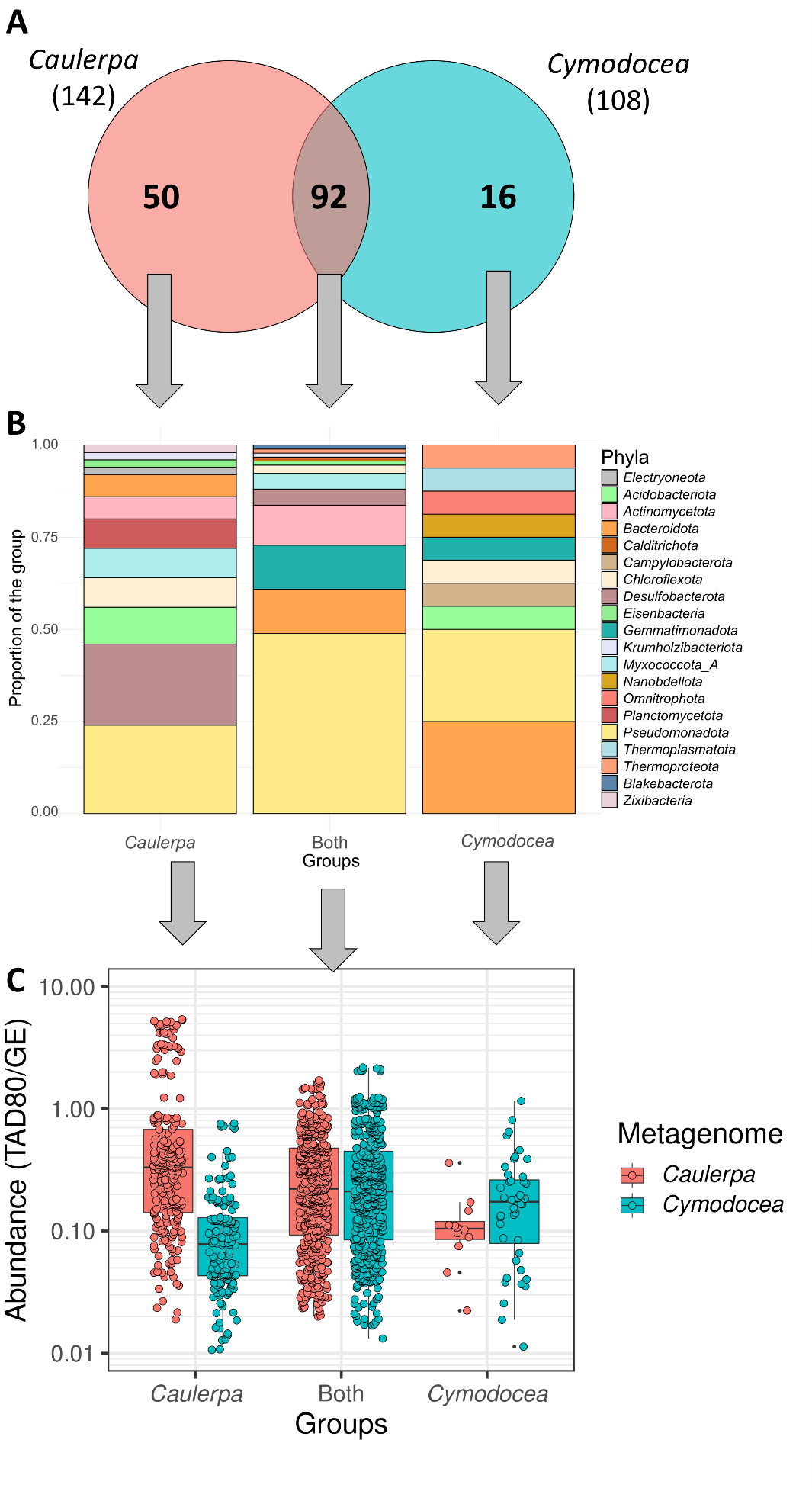

**Figure S3.** A) Venn diagram showing the MAGs included in the “*Caulerpa*,” “*Cymodocea*,” and “both” groups. B) Taxonomic classification of the MAGs included in each group, expressed as the percentage represented by each phylum within the group. C) Boxplot of the abundance of the MAGs included in each group as a function of metagenome type, *Caulerpa* or *Cymodocea*.

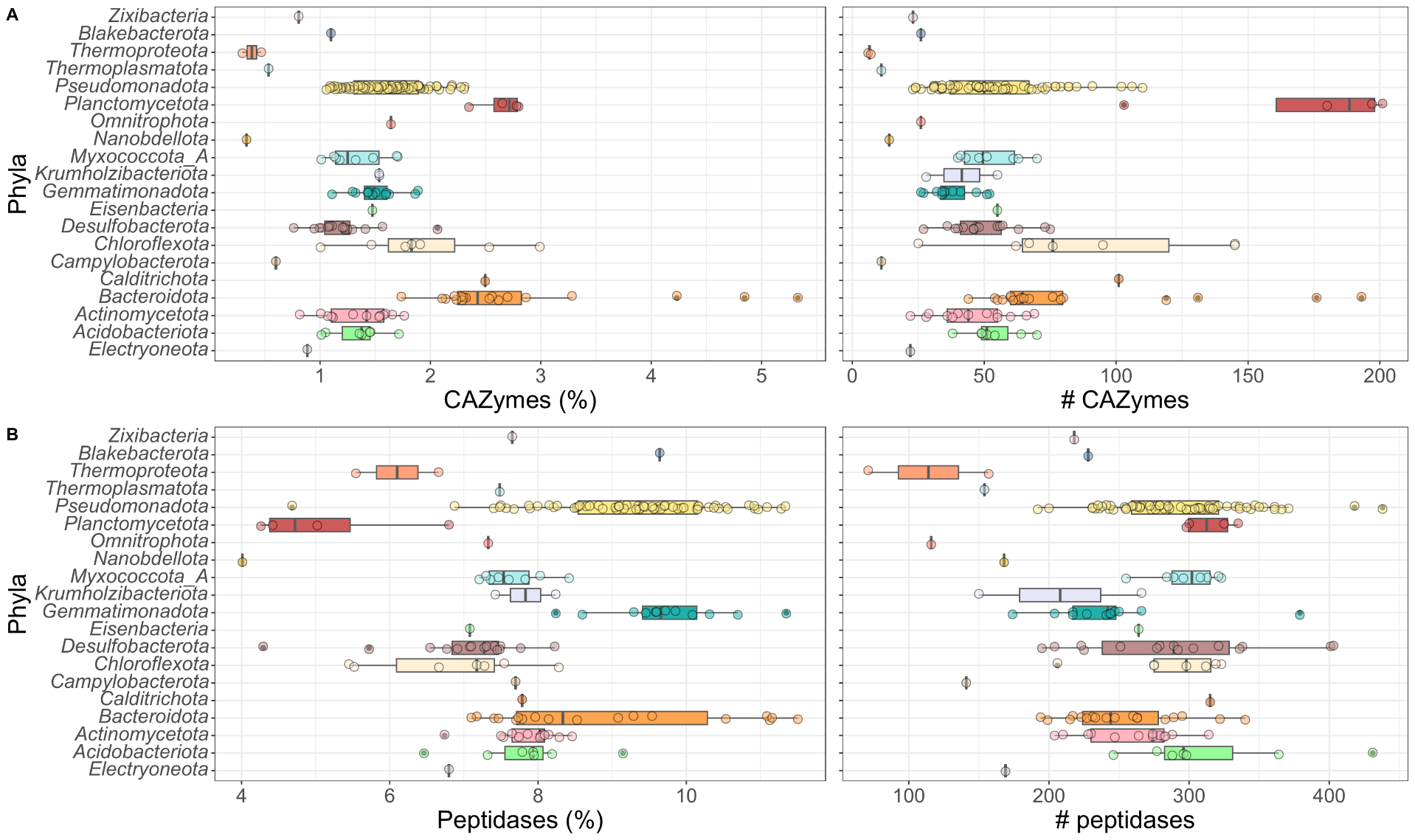

**Figure S4.** Abundance of putative carbohydrate-active enzymes (CAZyme) and peptidase encoding genes in MAGs recovered from the Mar Menor metagenomes. Boxplots show the proportion (left, x-axis) and absolute number (right, x-axis) of CAZyme (A) and peptidase (B) encoding genes across MAGs, grouped by phylum (y-axis).

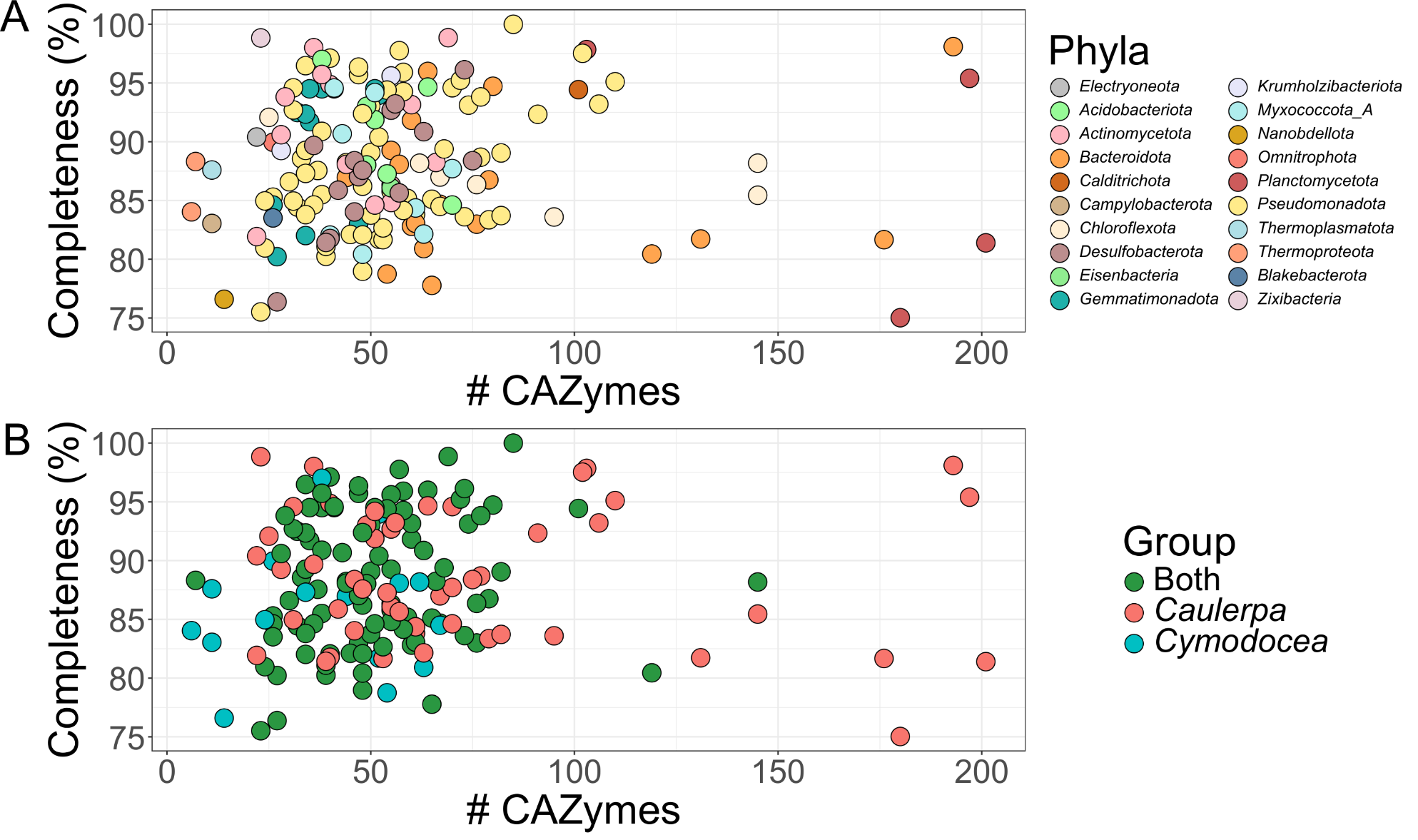

**Figure S5.** Scatter plot showing MAG completeness, as estimated by CheckM, on the y-axis versus the number of putative CAZy encoding genes on the x-axis. Point colors indicate the phylum (A) or the group (B) to which each MAG belongs.

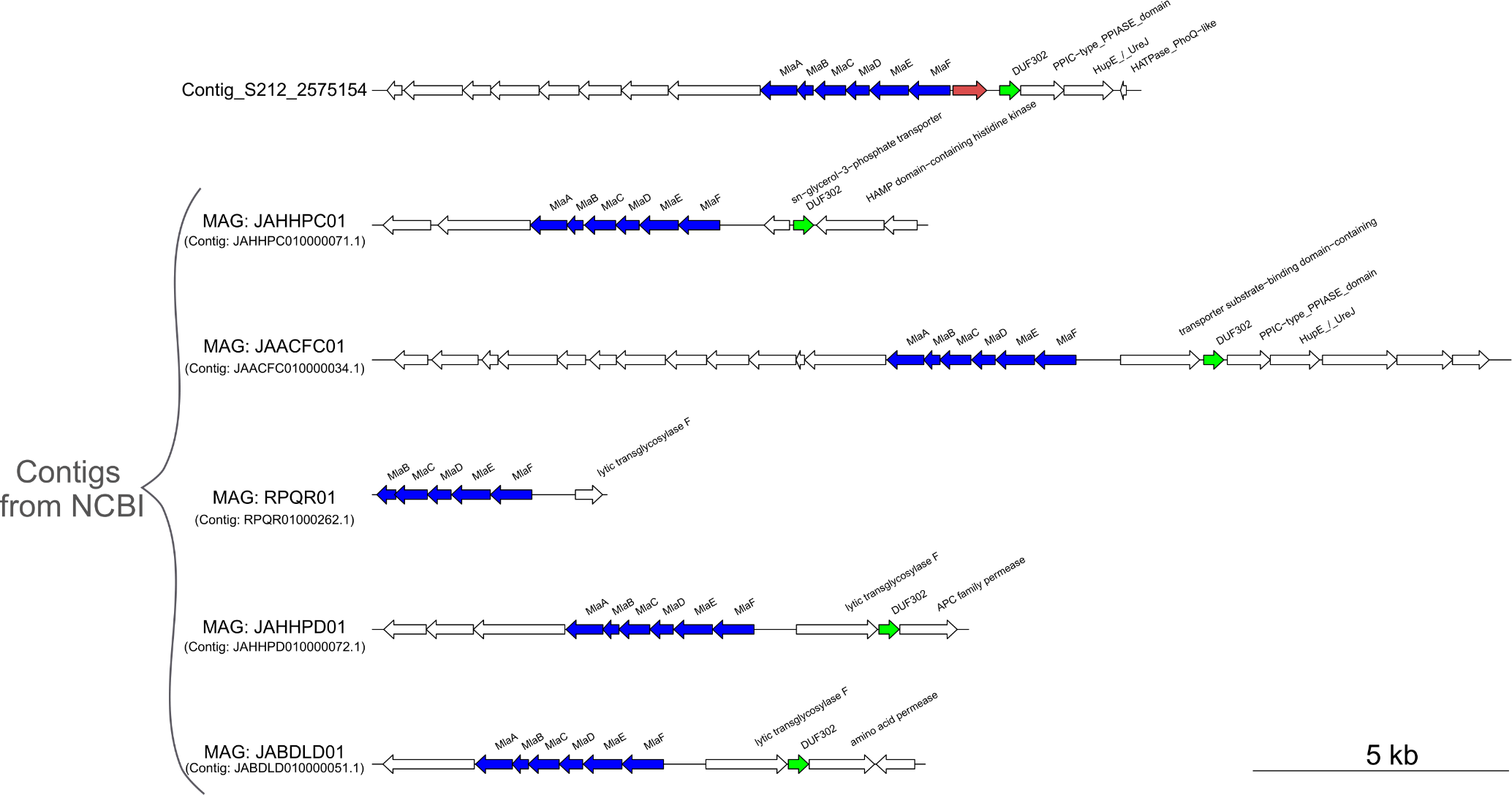

**Figure S6.** On top, gene organization in the contig encoding the highly expressed protein VHTgp (red gene) belonging to a *Gammaproteobacteria* MAG recovered from the Mar Menor metagenomes. Below, gene organization of five contigs from MAGs deposited in NCBI classified within the *Gammaproteobacteria*. Note the global distribution of these four contigs; MAG JAACFC01 was collected from marine sediments in China, MAG RPQR01 from a freshwater lake in northern United States and MAGs JAHHPD01 and JABDLD01 from coastal sediments in Australia.

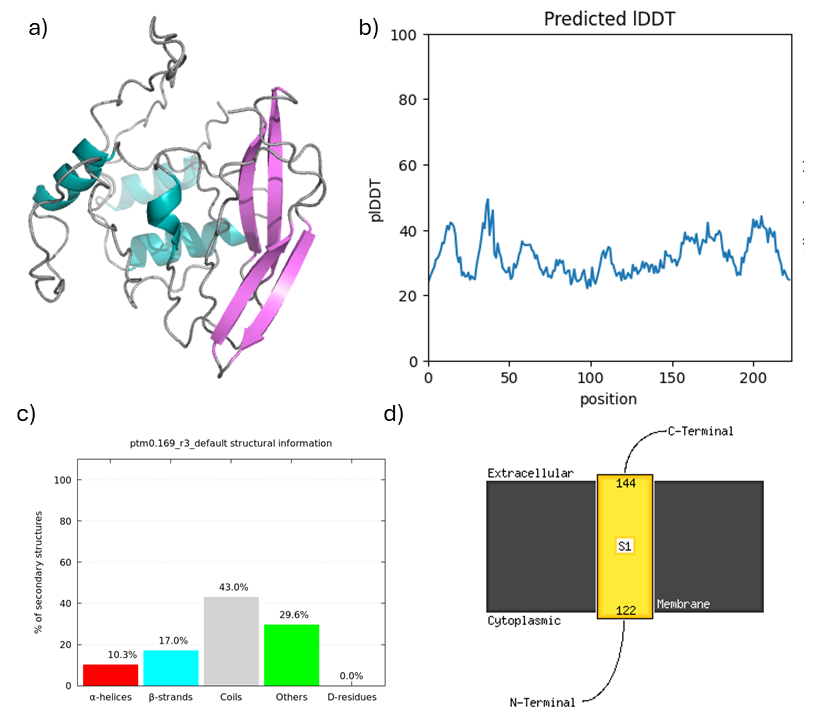

**Figure S7.** A) ESM Fold model of VHTgp shows α-helices (10%, light blue) and β-sheets (17%, pink) but mostly the structure is ill-defined. B) confidence score (pLDDT) across the protein is low between 20 – 40%. C) The proportions of secondary structures in the ESMFold model after calculating the predicted CD spectrum [1]. D) MEMSAT-SVM representation of the C-terminal of VHTgp spanning the membrane.

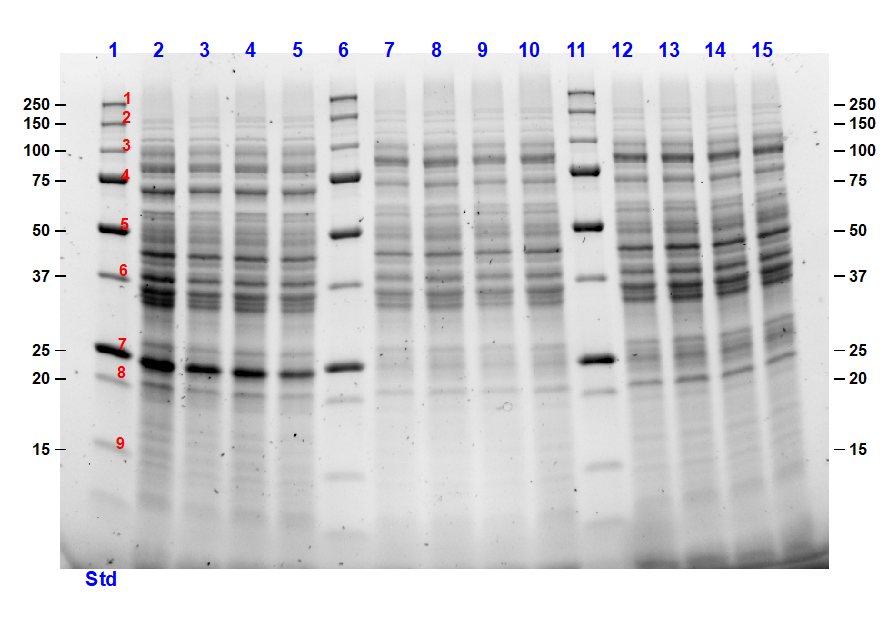

**Figure S8.** Whole cell lysates analyzed by SDS-PAGE of pET23a(VHTgp); lanes 2 – 5, pET28a(VHTgp); lanes 7 – 10, and empty pET28 control vector; lanes 12 –15.

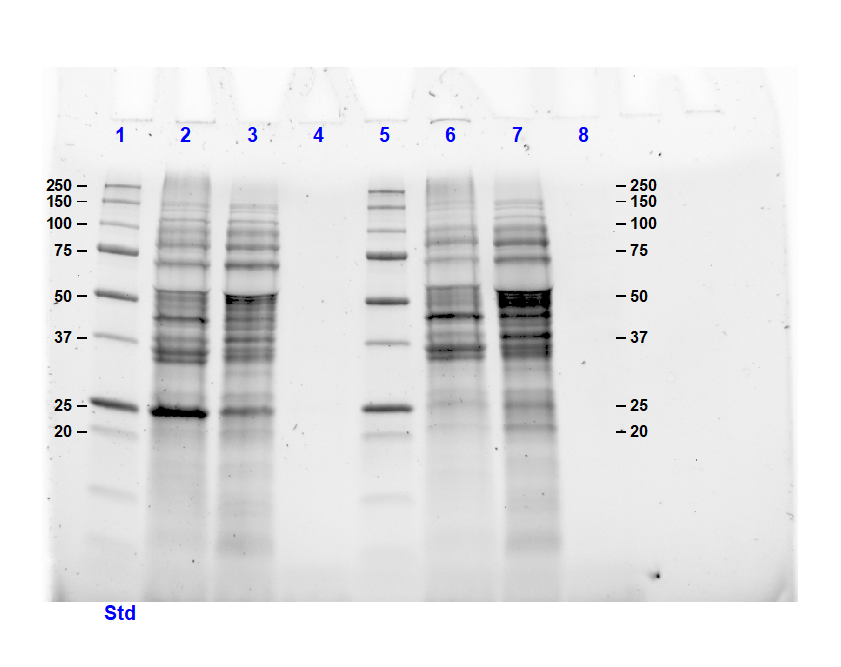

**Figure S9**. SDS-PAGE of insoluble fractions pET23a(VHTgp) (lane 2) and pET28a(VHTgp) (lane 6); clarified cell lysates of pET23a(VHTgp) (lane 3) and pET28a(VHTgp) (lane 7); elution fraction after IMAC purification of pET23a(VHTgp) (lane 4) and pET28a(VHTgp) (lane 8).

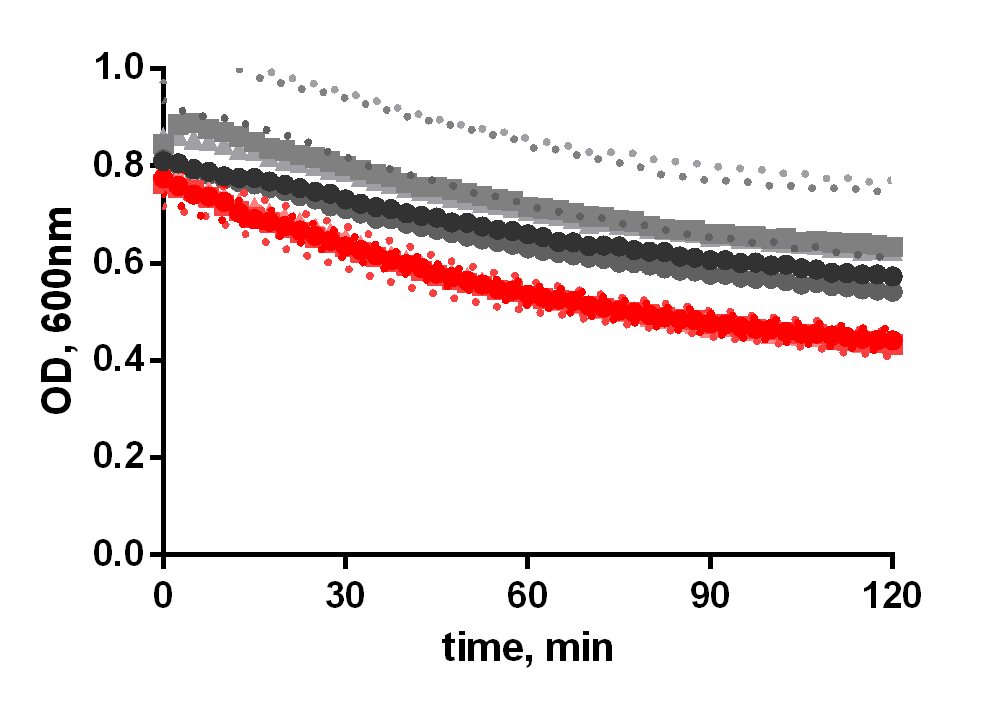

**Figure S10**. Turbidity Reduction Assay of clarified cell lysates VHTgp (red), empty pET28a (gray) and CEWL (black) after boiling for 10 minutes. Circle, square and triangle represent biological replicates, dotted lines represent the standard deviation of the average of three replicates. OD_600nm_=40 for all samples.
